# Short-term oxycodone exposure produces delayed and persistent gut microbiome disruption in mice

**DOI:** 10.64898/2026.05.26.727957

**Authors:** Junyi Tao, Daniel Gomez, Yaa F. Abu, Kristin Rojas, Sabita Roy

**Affiliations:** Department of Surgery, University of Miami Miller School of Medicine, Miami, FL, 33136, USA

## Abstract

The gut microbiome is a critical part of host homeostasis, yet its resilience following opioid exposure remains poorly understood. While opioid-induced short-term dysbiosis is well documented, the long-term recovery dynamics following oxycodone remain unclear. This study characterized the temporal dynamics of the fecal microbiota in male C57BL/6J mice following a brief 3-day oxycodone regimen (5mg/kg, BID). 16S rRNA gene sequencing was performed at baseline, day 3, 10, 17, and 70. While acute post-treatment phases (day 3 to 10) showed subtle taxonomic shifts in *Clostridium_sensu_stricto_1* and *Romboutsia*, significant community disruption emerged later. By day 17, beta diversity significantly differed from saline controls (P =0.002). At day 70, both alpha diversity (p=0.02) and beta diversity (P=0.007) remained significantly altered, characterized by enriched *Akkermansia* and *Marvinbryantia* alongside depleted *Eubacterium_xylanophilum*. These findings demonstrate that even brief oxycodone exposure triggers persistent, non-recovering dysbiosis that became detectable only after treatment cessation and persisted through day 70. This suggests that the window for microbiome recovery exceeds two months in mice (equivalent to several human years), highlighting a potential long-term risk for patients prescribed short-term opioid courses.

**Importance:** Short-term opioid exposure is generally assumed to cause only transient disruption of the gut microbiome. However, the duration of microbiome recovery following clinically relevant opioid treatment remains poorly defined. In this study, we show that a brief three-day course of oxycodone in mice resulted in delayed and persistent alterations in gut microbial community structure that remained detectable for at least 70 days after treatment cessation. Notably, significant divergence in microbial composition emerged weeks after exposure rather than immediately following treatment, suggesting that short-term opioid use may initiate longer-lasting remodeling of the gut microbiome than previously appreciated. These findings highlight the importance of considering extended recovery timelines when evaluating the microbiological consequences of opioid exposure.

## OBSERVATION

The gut microbiome, a complex community of bacteria and fungi, plays a critical role in maintaining host homeostasis. However, its stability is significantly compromised by exogenous stressors such as opioid analgesics. Extensive research has established that both acute and chronic opioid administration induce profound gut dysbiosis, characterized by altered microbial diversity and a shift toward pathogenic taxonomic profiles (1). While many studies utilize morphine as a model opioid, oxycodone represents a more clinically relevant opioid due to its high rates of prescription for acute, perioperative, and chronic pain management (2). Despite its clinical prevalence, a significant gap remains in the current literature regarding the resilience of the microbiome following drug cessation. Specifically, it remains unclear whether the gut microbiota eventually reverts to its baseline state or if oxycodone-induced shifts persist as a long-term disruption of the microbial landscape. To address this, the current study aims to characterize the temporal dynamics of oxycodone-induced dysbiosis and determine the duration required for the microbiome to recover its original state in a mouse model.

Male C57BL/6J mice (12□weeks old) were obtained from The Jackson Laboratory. Mice were randomly assigned to experimental groups and provided with the same diet and living conditions, with food and water available *ad libitum*. Twenty mice were randomly assigned to either an oxycodone treatment group (oxy) or a saline control group (sal) (n=10 per group). Animals received subcutaneous injections of either oxycodone (5 mg/kg) or an equivalent volume of saline twice daily for three consecutive days (Figure S1).

Fresh fecal pellets were collected from individual mice at five distinct time points: day 0 (baseline), day 3 (completion of treatment), and days 10, 17, and 70 (representing 1-week, 2-week, and 2-month post-injection intervals, respectively) (Figure S1). All samples were immediately flash-frozen and stored at −80°C until further processing. DNA was isolated from fecal samples using a DNeasy 96 PowerSoil Pro QIAcube HT Kit with a QIAcube HT liquid-handling machine.

During DNA extraction, two extraction controls in each batch were included to account for and identify potential contamination. Sequencing was performed by the University of Minnesota Genomics Center (UMGC). The hypervariable V4 region of the 16S rRNA gene was PCR amplified using the forward primer 515F (GTGCCAGCMGCCGCGGTAA), reverse primer 806R (GGACTACHVGGGTWTCTAAT), Illumina adaptors, and molecular barcodes to produce 420-bp amplicons. Amplicons were sequenced with the Illumina MiSeq version 3 platform, generating 300-bp paired end reads. The FASTQ files were clustered into amplicon sequence variants (ASVs) using the DADA2 package (version 1.26.0) (3). Taxonomic assignment of ASVs was assigned to the species level using a naïve Bayesian classifier (4) implemented in DADA2 with the SILVA reference database (release 138.1) (5). ASV and taxonomy table can be accessed in Table S1 for all samples. ASV and taxonomy tables were imported into MicrobiomeAnalyst (6) for generating alpha and beta diversity plots, taxonomy bar plots and linear discriminant analysis effect size (LEfSe) (7) plot. The threshold on the logarithmic LDA score for discriminative features was set to 2. The Benjamini-Hochberg method was used for controlling the false-discovery rate (q value). The cutoff for q value was set to 0.1 for LEfSe analysis. Mann– Whitney test was used to detect if α-diversity differed across treatments. Permutational multivariate analysis of variance (PERMANOVA) was used to detect if β-diversity differed across treatments.

For this study, longitudinal fecal sampling was performed from the same animals across all timepoints (Figure S1). To account for the confounding effects of coprophagy and the resulting homogenization of the gut microbiota within shared housing, fecal samples were analyzed as group-level cross-sectional profiles at each timepoint. First, we compared the diversity and composition difference between the oxy and sal group to see if the starting microbiome is similar between groups. When β diversity was measured Bray-Curtis distances and visualized with principal-coordinate analysis (PCoA) plots, fecal samples from baseline saline mice (sal_day0) clustered together with samples from pre-treated oxycodone mice (oxy_day0) (p = 0.61) (Figure S2A). The α-diversity was measured by Shannon index which accounted for both sample richness and evenness. Similarly, the α-diversity was not statistically different between sal_day0 and oxy_day0 groups (p = 0.71) (Figure S2B). LEfSe analysis did not identify any significantly different feature between sal_day0 and oxy_day0 groups. Taken together, both groups had similar microbiome before the treatment.

Following the three-day treatment period, alterations in microbial composition were detectable as early as day 3. On day 3, when β diversity was measured using Bray-Curtis distances and visualized with PCoA plots, fecal samples from saline-treated mice (sal_day3) clustered together with samples from oxycodone-treated mice (oxy_day3) (p = 0.13) (Figure 1A). The α-diversity measured by Shannon index was not different between sal_day3 and oxy_day3 groups (p = 0.9) (Figure 1C). LefSe analysis was also performed between the groups to determine the bacterial genera that were differentially enriched on day 3. LEfSe analysis demonstrated that genera *Clostridium_sensu_stricto_1, Romboutsia*, and *UCG-009* were more abundant in oxy_day3 group (Figure 1E). No bacteria genera were more abundant in sal_day3 group.

**Figure 1.**
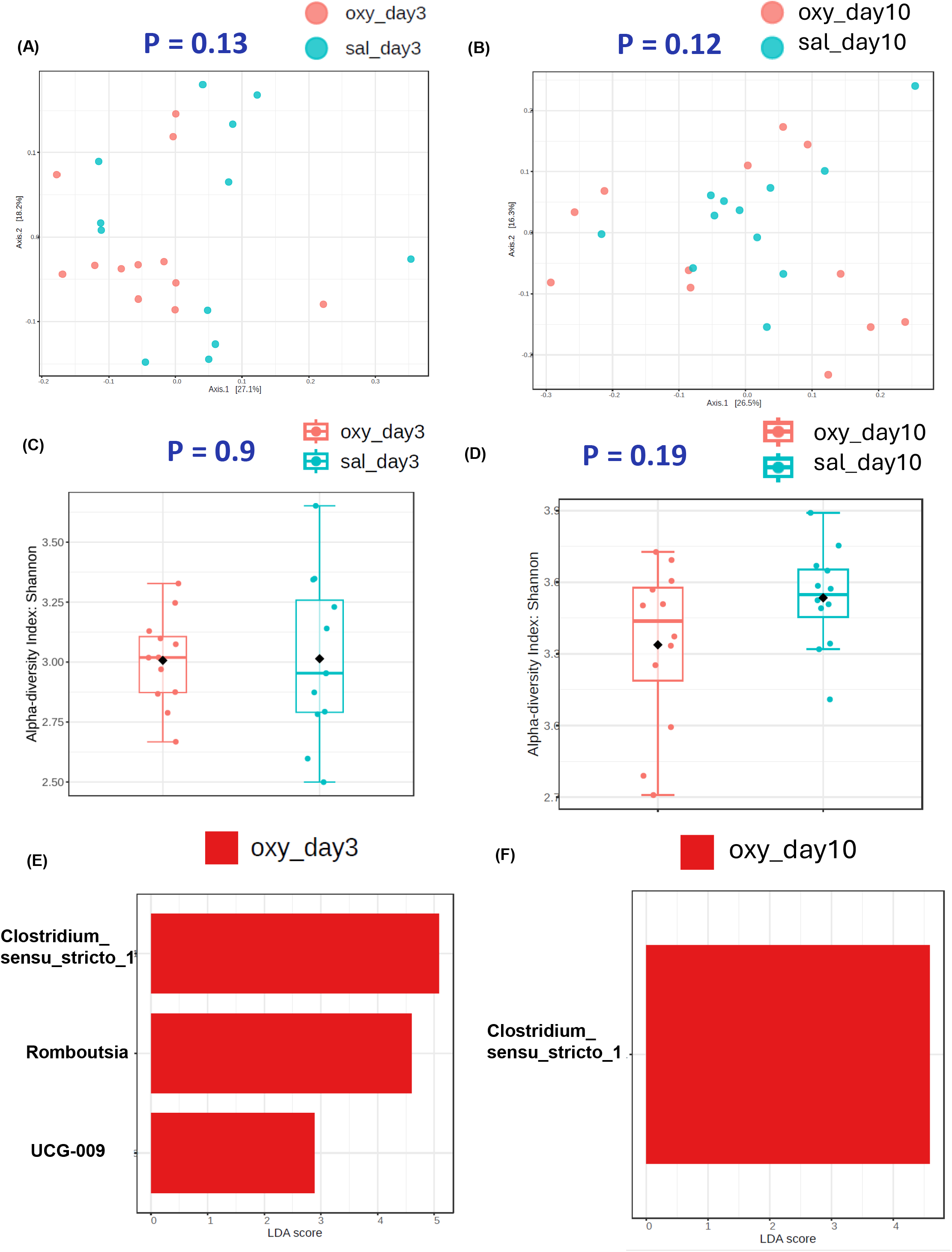
Diversity and composition analysis of the fecal microbiome samples on day 3 and day 10. Samples are grouped by saline group on day 3(sal_day3) (n□=□12) oxycodone group on day 3 (oxy_Day3) (n□=□12), saline group on day 10(sal_day10) (n□=□12) and oxycodone group on day 10 (oxy_Day10) (n□=□12). (A). PCoA plot of Bray-Curtis distance (metrics of β - diversity) between groups on day 3. (B). PCoA plot of Bray-Curtis distance (metrics of β -diversity) between groups on day 10. (C) Shannon index (metrics of α-diversity) between groups on day 3. (D) Shannon index (metrics of α-diversity) between groups on day10. (E). LefSe analysis of top discriminative bacteria genera between groups on day 3. (F). LefSe analysis of top discriminative bacteria genera between groups on day 10. UCG is short for uncultured genus-level group.

Altered microbiome composition was maintained through day 10. On day 10, when β diversity was measured using Bray-Curtis distances and visualized with PCoA plots, fecal samples from saline-treated mice (sal_day10) clustered together with samples from oxycodone-treated mice (oxy_day10) (p = 0.12) (Figure 1B). The α-diversity measured by Shannon index was not different between sal_day10 and oxy_day10 groups (p = 0.19) (Figure 1D). LefSe analysis was also performed between the groups to determine the bacterial genera that were differentially enriched on day 10. The bar plot of the LDA score demonstrated that one of the genera that were more abundant in oxycodone-treated group at day 3, *Clostridium_sensu_stricto_1*, were also more abundant in oxy_day10 group (Figure 1F). No bacteria genera were more abundant in sal_day10 group. *Clostridium_sensu_stricto_1* are strictly anaerobic, and have the ability to metabolize various compounds such as carbohydrates, amino acids, alcohols and purines (8). Butyric acid is a ‘genus specific’ product of fermentation (9) for *Clostridium_sensu_stricto_1*. Interestingly, this genus has been significantly correlated with the expression of inflammation-related genes, including *REG3G, CCL8*, and *IDO1*(10). Genus *Romboutsia* was higher in oxy group on day 3. *Romboutsia* species are flexible anaerobes that are adapted to a nutrient-rich environment where carbohydrates, amino acids, and vitamins are abundantly available (11). In sum, a few bacteria genera were altered by the 3-day oxycodone treatment, and the alteration lasted for at least for 1 week from day 3 to day 10. However, the treatment did not alter the overall diversity metrics during the first week post treatment period.

By day 17, significant alterations in both microbial diversity and composition emerged. On day 17, when β diversity was measured using Bray-Curtis distances and visualized with PCoA plots, fecal samples from saline-treated mice (sal_day17) significantly clustered separately with samples from oxycodone-treated mice (oxy_day17) (p = 0.002) (Figure 2A). However, the α-diversity measured by Shannon index showed no difference between sal_day17 and oxy_day17 groups (p = 0.24) (Figure 2C). LefSe analysis was also performed between the groups to determine the bacterial genera that were differentially enriched on day 17. The bar plot of the LDA score demonstrated that genera *Eubacterium_xylanophilum* and ASF 356 were more abundant in oxy_day17 group (Figure 2E). Genera *Monoglobus*, and *Dubosiella* were more abundant in sal_day17 group (Figure 2E).

**Figure 2.**
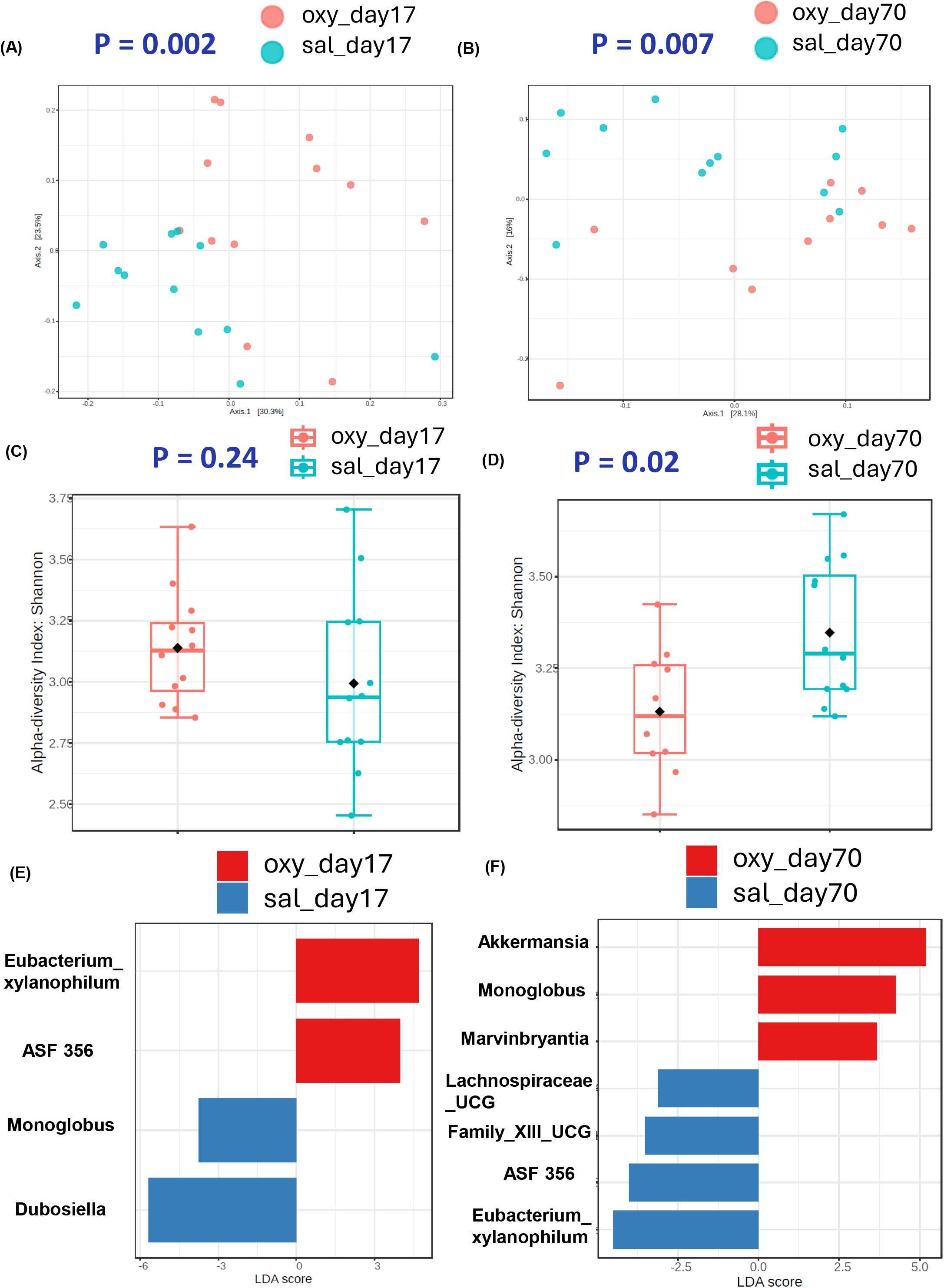
Diversity and composition analysis of the fecal microbiome samples on day 17 and day 70. Samples are grouped by saline group on day 17(sal_day17) (n□=□12) oxycodone group on day 17 (oxy_Day17) (n□=□11), saline group on day 70(sal_day70) (n□=□12) and oxycodone group on day 70 (oxy_Day70) (n□=□10). (A). PCoA plot of Bray-Curtis distance (metrics of β - diversity) between groups on day 17. (B). PCoA plot of Bray-Curtis distance (metrics of β -diversity) between groups on day 70. (C) Shannon index (metrics of α-diversity) between groups on day 17. (D) Shannon index (metrics of α-diversity) between groups on day70. (E). LefSe analysis of top discriminative bacteria genera between groups on day 17. (F). LefSe analysis of top discriminative bacteria genera between groups on day 70. UCG is short for uncultured genus-level group.

The alteration in microbiome diversity and composition persisted on day 70. When β diversity was measured using Bray-Curtis distances and visualized with PCoA plots, fecal samples from saline-treated mice (sal_day70) significantly clustered separately with samples from oxycodone-treated mice (oxy_day70) (p = 0.007) (Figure 2B). Notably, the α-diversity measured by Shannon index was significantly lower in oxy_day70 group than sal_day70 group (p = 0.02) (Figure 2D). LefSe analysis was also performed between the groups to determine the bacterial genera that were differentially enriched on day 70. The bar plot of the LDA score demonstrated that genera *Eubacterium_xylanophilum*, ASF 356, *Lachnospiraceae_UCG*, and *Family_XIII_UCG* were more abundant in sal_day70 group (Figure 2F).

Genera *Akkermansia, Monoglobus*, and *Marvinbryantia* were more abundant in oxy_day70 group (Figure 2F). Persistent depletion of taxa such as *Eubacterium xylanophilum* and enrichment of *Akkermansia* suggest altered mucin utilization and short-chain fatty acid metabolism as potential downstream consequences of transient opioid exposure. Interestingly, *Eubacterium xylanophilum* and ASF 356 were more abundant in oxy group on day 17 but were less abundant in oxy group on day 70. In a preclinical study, *Eubacterium xylanophilum group* was statistically more abundant in the mice group that was fed rice bran supplement (12). Strain ASF 356 is most closely related to *Clostridium propionicum* (13). *Clostridium* ASF356 produced the common fermentation end products acetate, propionate, succinate, and butyrate (14). Conversely, the genus *Monoglobus* has the opposite trend. It is more abundant in saline group on day 17 but less abundant in saline group on day 70. *M. pectinilyticus* is capable of fermenting pectins as well as carbohydrates that are constituents of pectins and hemicellulose, such as galacturonic acid, xylose, and arabinose (15). In sum, brief 3-day treatment of oxycodone was enough to induce long-lasting dysbiosis in mice to at least 70 days.

Several studies have reported that chronic oxycodone exposure has no significant impact on the gut microbiome. For instance, in a study that was conducted on male rats, oxycodone was administered subcutaneously 2 x a day using incremental increases for 12 days. Fecal samples were collected after last day of treatment, and no difference was observed in beta diversity, alpha diversity or Firmicutes/Bacteroidetes ratio among the treatment groups (16). In another study with rat model, similar results were reported where oxycodone dependence was initiated by injecting the rats 2 times a day for 5 days. The PCoA plot measured by Bray–Curtis distance indicated that the saline and oxycodone groups clustered together on day 12. However, other clinical and pre-clinical studies reported that oxycodone treatment had effects on microbiome composition. Consistent with our findings, a clinical cohort of 25 patients with moderate to severe cancer pain, abundance of *Lactobacillus, Anaerostipes Megamonas, Monoglobus*, and *Rikenellaceae_ RC9_gut_group* were significantly changed between the oxycodone group and the control group (17). There were no significant differences in the Chao 1 and Shannon diversity indexes among the control and oxycodone patients. Additionally, no significant difference in beta diversity among the control, oxycodone-sensitive and oxycodone-tolerant groups(17). Furthermore, Lyu et al reported that *in utero* exposure to oxycodone for 2 weeks in mouse model can lead to long tern effects on the gut microbiota of the offspring when examined at adulthood (18). In their study, 120-day-old male offspring showed greater oxy vs. sal segregation than female offspring, echoing the significant long-term shifts we observed in our male cohort at day 70. In another mice study where animals in the oxycodone group were given oxycodone injections for 5 weeks at escalating doses, showed distinct separation from control group in PCoA plot measured by both Jaccard and Bray-curtis distances (19). Moreover, α-diversity measured by CHAO1 was significantly lower in the control group when compared to those treated with morphine or oxycodone.

The discrepancies between our findings and previous research likely stem from several factors. Notably, the two studies reporting no significant effects of oxycodone utilized rat models, whereas studies indicating positive effects focused on humans or mice. The timing of sampling also appears to be critical to detecting the delayed evolution of dysbiosis, even in a controlled environment. The majority of the studies collect the samples the same day or the next day after the last day of oxycodone treatment, and their results were similar to our results on day 3. In contrast, we continued to collect samples after the cessation of oxycodone treatment, for up to 70 days, which represents a prolonged recovery interval relative to murine lifespan (approximately 8.6 years in human) (20). Previous investigators might have detected a difference in beta diversity if they followed up their samples for months instead of days. A limitation of our study included limiting the test subjects to male, instead of including female mice. This could have been particularly helpful when correlating our findings to those of Lyu, et al (18).

Our findings indicate that microbiome recovery following short-term oxycodone exposure is substantially delayed and remains incomplete for at least 70 days after treatment cessation. In summary, a brief three-day oxycodone exposure produced delayed and persistent alterations in gut microbial community structure that remained evident for at least 70 days after treatment cessation. These findings suggest that short-term exposure to clinically relevant opioids may induce longer-lasting microbiome disruption than previously appreciated and highlight the importance of extended recovery timelines when evaluating opioid-associated microbiome effects.

## Ethics

Animal studies were conducted in accordance with protocols set forth by the University of Miami Institutional Animal Care and Use Committee (IACUC) and in compliance with rules and guidelines described by the National Institute of Health Guide for the Care and Use of Laboratory Animals.

## Data Availability

The raw fastq files have been uploaded to NCBI Bioproject with accession number: PRJNA1443279. The data structure can be viewed via reviewer link https://dataview.ncbi.nlm.nih.gov/object/PRJNA1443279?reviewer=6ehahh7qj2vssf66scgss4hcp4.

## ACKNOWLEDGMENTS

This work was supported by National Institutes of Health, Grant/Award Numbers: R01 DA043252 (Roy), R01 DA044582 (Roy), R01 DA047089 (Roy), R01 DA050542 (Roy), P30 CA240139_supplement5 (Rojas).

## Suplemental Figures

**Figure S1.**
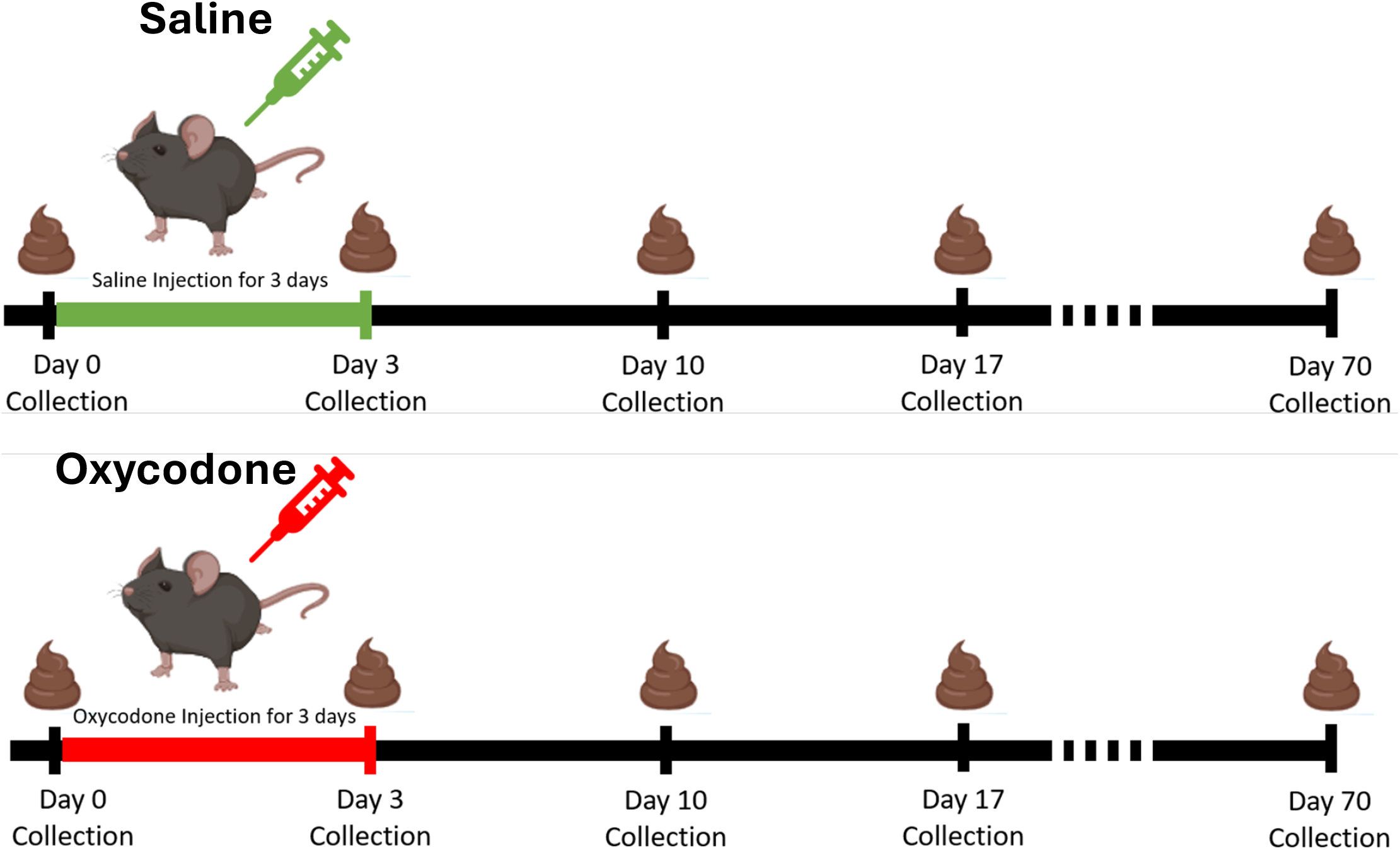
Schematic of oxycodone treatment regime in the mouse model.

**Figure S2.**
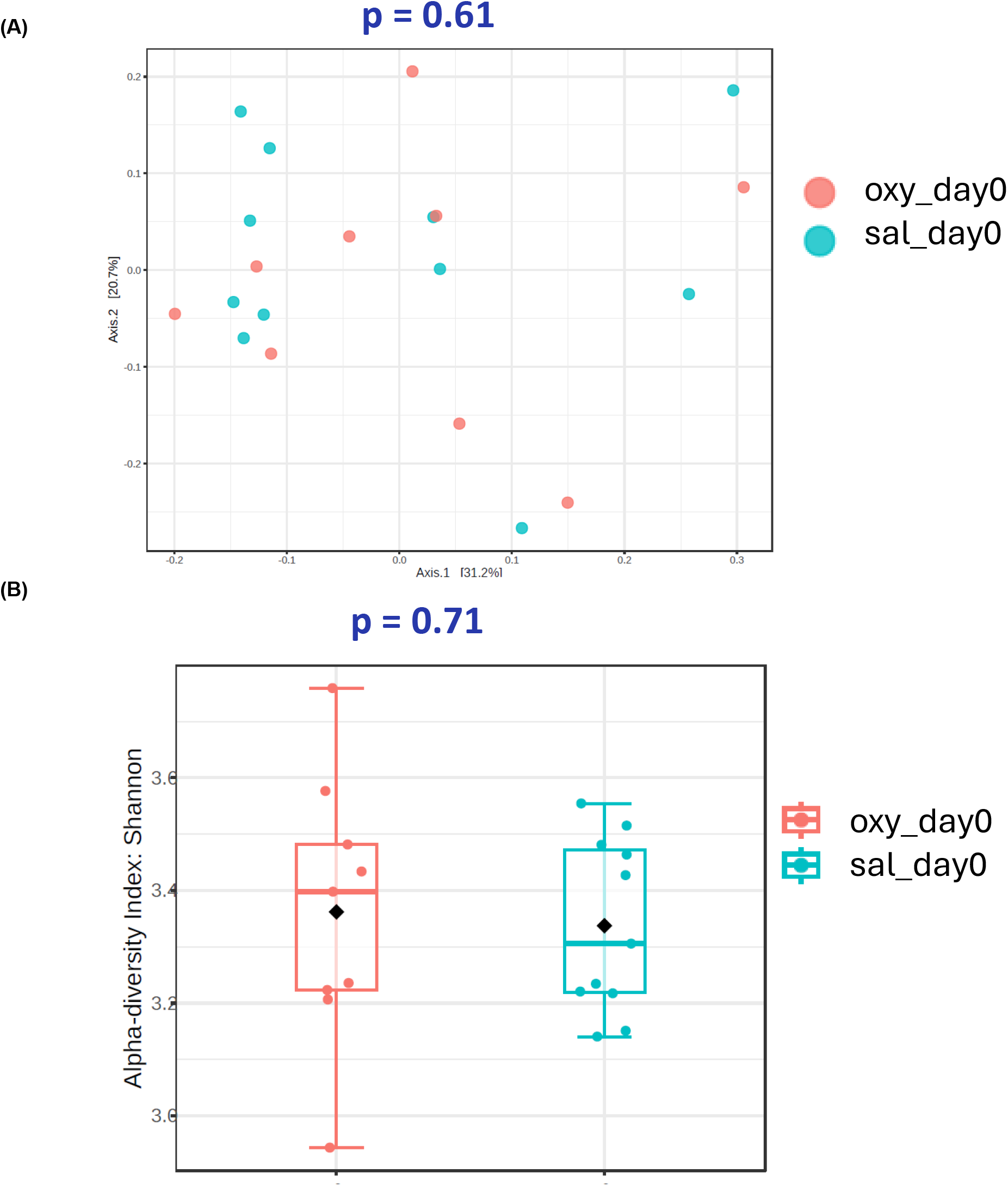
Diversity analysis of the fecal microbiome samples on day 0. Samples are grouped by saline pre-treated group (sal_day0) (n = 11) and oxycodone pre-treated group (oxy_Day0) (n = 9). (A). PCoA plot of Bray-Curtis distance (metrics of β -diversity) between sal_day0 and oxy_day0. (B) Shannon index (metrics of α-diversity) at day 0. Error bars represent SEM.

